# TRGdb: a universal resource for exploration of taxonomically restricted genes in bacteria

**DOI:** 10.1101/2023.02.17.528947

**Authors:** Andrzej Zielezinski, Wojciech Dobrychlop, Wojciech M. Karlowski

## Abstract

The TRGdb database is a resource dedicated to taxonomically restricted genes (TRGs) in bacteria. It provides a comprehensive collection of genes that are specific to different genera and species, according to the latest release of bacterial taxonomy. The user interface allows for easy browsing and searching as well as sequence similarity exploration. The website also provides information on each TRG protein sequence, including its level of disorder, complexity, and tendency to aggregate. TRGdb is a valuable resource for gaining a deeper understanding of the TRGs-associated, unique features and characteristics of bacterial organisms. The TRGdb resource is freely accessible through www.combio.pl/trgdb.

## Introduction

Taxonomically Restricted Genes (TRGs) are genes that are present only in a particular taxonomic unit and have no traceable evolutionary history [1]. As the TR genes have no homology to genes from other organisms, they are fundamentally important for the study of the emergence of organismal unique traits such as morphological diversity, metabolic innovation, and pathogenicity [2]. Additionally, TRGs serve as taxon-specific diagnostic targets that can be used to reconstruct the evolutionary history of specific phylogenetically-related groups of organisms [3,4]. TRGs that are present at the species and genus levels garner special interest, because they are expected to perform a role in defining exclusive ecological adaptations of organisms to particular niches [4,5].

With whole-genome sequencing further facilitated by next-generation technologies, species- and genus-specific TR genes continue to be discovered in every newly sequenced genome across prokaryotes [1,5–12], eukaryotes [13–17], and viruses [18,19]. Although eukaryotic TRGs are important for understanding the diversity and complexity of organisms, studying bacterial TRGs is crucial for gaining insight into the properties and evolution of the biotechnologically, biomedically, and scientifically important microorganisms [1]. Paradoxically, although information on bacterial TRGs is essential for advancing our understanding of bacterial biology and evolution, it is not readily accessible. Two pioneering databases of species-specific TRGs in bacteria, OrphanMine [5] and ORFanage [20], have been unavailable for many years now, which hinders attempts at a systematic analysis of taxonomically restricted genes in the domain of bacteria.

To reconcile the deficiency of the current surveys of TRGs, we present a comprehensive database of species-and genus-specific taxonomically restricted genes in bacteria. By applying the TRG identification scheme from our recent work [21], we analyzed available genomes of more than 62 thousand bacterial species represented by almost 200 million proteins. We supply a user-friendly interface for the results of the predictions in the context of the most up-to-date bacterial taxonomy, allowing users to browse and search for TRGs across different taxonomic units of bacteria. We also provide information on every TRG protein, including its sequence properties (e.g., level of disorder and complexity, and tendency to aggregate), affiliation to the TR gene family, and links to external databases.

## Methods

### Data sources

The sequence data, information about the taxonomic classification, and a phylogenetic tree of bacteria were obtained from the Genome Taxonomy Database (GTDB) release 07-RS207 (April 2022) [22]. The data set includes 62,291 representative genomes, with one genome per species. The GTDB database provides high-quality bacterial taxonomy based on phylogenetic analysis and carefully selects the best representative genome for each species. The representative genomes in GTDB have the highest assembly quality, the least amount of contamination in the sequence, and the most complete set of genes.

### Identification of TRGs

Our TRG identification procedure [21] includes several steps that allow high-quality TR gene predictions. However, this procedure can currently be applied only to a limited-sized data set (e.g., a single genus). Therefore, the approach employed in the construction of TRGdb was simplified and limited to three progressive steps. First, we used DIAMOND v2.0.15 [23] to perform an all-versus-all comparison between protein sequences (*n* = 193,808,833) from 62,291 bacterial species. Then, we removed from further analysis query proteins that had highly similar (homologous) sequence matches (*E*-value ≤ 10^−3^) belonging to bacterial species outside the genus of the query species. This step allowed us to reduce the number of candidate TRGs by approximately 90%. The remaining sequences were classified as the candidate TR genes at the genus level. Next, we verified the TRG candidates by querying them with BLAST+ v2.13.1 [24]. The number of reported hits per query (*-max_target_seqs*) was adjusted to accommodate for the number of all tested species in a given genus. Query that did not show significant similarity (*E-value* ≤ 10^−3^) to any sequences from organisms outside the tested genus were identified as genus-specific TR genes. Finally, we extracted a list of bacterial species-specific sequences from the obtained TR gene list. These genes were defined as protein sequences that did not have homologs outside the query species and the genus of query species encompassed at least two species according to the GTDB taxonomy.

### Sequence properties calculations

To determine the properties of the TRG protein sequences, we followed the methodology described in [21]. We used IUPred2 [25] to calculate the disorder score of the TRG proteins, which reflects the extent of the intrinsic disorder in the proteins. This score was calculated by dividing the number of residues with a disorder score above 0.5 by the total length of the protein sequence. We used TANGO v2.3.1 [26] to determine the TRG protein’s average aggregation, which was presented as the frequency of potential aggregating segments defined as hexapeptides with an aggregation score above 5% of all amino acid residues. Finally, we used SciPy v1.21.4 [27] to calculate the sequence complexity of the TRG proteins, which was assessed as Shannon entropy.

### TRG protein families

We classified the TRG proteins into families, similarly to how protein families are established in the Pfam-B database [28]. Accordingly, we used <u>MMseqs2</u> v14-7e284 [29] with the cluster option. We set the maximum *E*-value at 10^−3^ and used the bidirectional coverage mode (*–cov-mode 0*) with a minimum coverage of 0.8 (*-c 0*.*8*).

### Word tags of protein annotations

We divided each line of the description from the NCBI protein record into individual words and manually removed common and meaningless terms (such as numbers and punctuation characters). Only words with at least 10 appearances were considered. The word cloud was created using WordArt (wordart.com), and the placement and color of word tags were adjusted manually using Inkscape v1.2.

### Calculation of Isolation index of Organisms (IIO)

To determine the degree of phylogenetic isolation of individual bacterial taxa, we calculated the Isolation Index of Organism (IIO) [6]. In our study, the IIO parameter was calculated based on the distance in the phylogenetic tree of bacteria in the GTDB database.

### Database interface and Application Programming Interface (API)

The TRGdb web interface was developed in HTML5 with the Bootstrap framework (v5.3), JavaScript, and Highcharts.js (v10.3.3). The database querying system was developed in Django (v4.0.0), Django REST framework (v3.13.1), and Python (v3.9.5) using the SQLite database as a data management system.

## Results

### Abundance and distribution of TRGs in Bacteria

We searched for the TR genes, conserved at the genus or species levels, in genomes of 62,291 bacterial species by developing the progressive TRG identification pipeline (see Methods) that accounts for up-to-date taxonomy of bacteria provided by the Genome Taxonomy Database (GTDB) (see Methods). We found almost 8.5 million TRG proteins, representing approximately 4.4% of all proteins in bacteria (**Table 1**). Of those proteins, 2.1% were found exclusively in genomes of a single species (referred to as species-specific TRGs) and 2.3% were present in the genomes of different species within the same genus (referred to as genus-specific TRGs). Almost every tested bacterial species (99.98%; **Table 1**) contains at least one TR gene at the species or genus level. On average, genus- and species-specific genes account for 4.6% of all genes in each genome, which is in accordance with early estimates of TRGs in bacterial species (1-16%) [1].

**Table 1.** Summary of the TRG identification results.

| Data type | Number |
| --- | --- |
| Bacteria species in total | 62,291 |
| Bacteria proteins in total | 193,808,833 |
| TRG proteins | 8,489,929 (4.38%) |
| Genus-specific TRG proteins | 4,444,919 (2.29%) |
| Species-specific TRG proteins | 4,045,010 (2.09%) |
| Bacteria species with TRG(s) | 62,279 (99.98%) |

To estimate the extent to which the separation of a species in a taxonomy tree of bacteria affects its TR gene count, we calculated an isolation index of organism (IIO), which represents a distance between a certain species or genus from the nearest other species or genus in a phylogenetic tree of bacteria (see Methods). We observed a moderate positive correlation (Pearson’s *r* = 0.4, *P* ≈ 0) between the number of TRGs and the IIO distance for the genus-level TRGs **(Figure 1a**), indicating a minor influence (*r*^*2*^ = 16%) of available genomic data (not sufficient taxonomic saturation) on the TRG content. However, the number of species-level TR genes does not seem to be impacted by the IIO distance (*r* = -0.02; **Figure 1b**).

**Figure 1.**
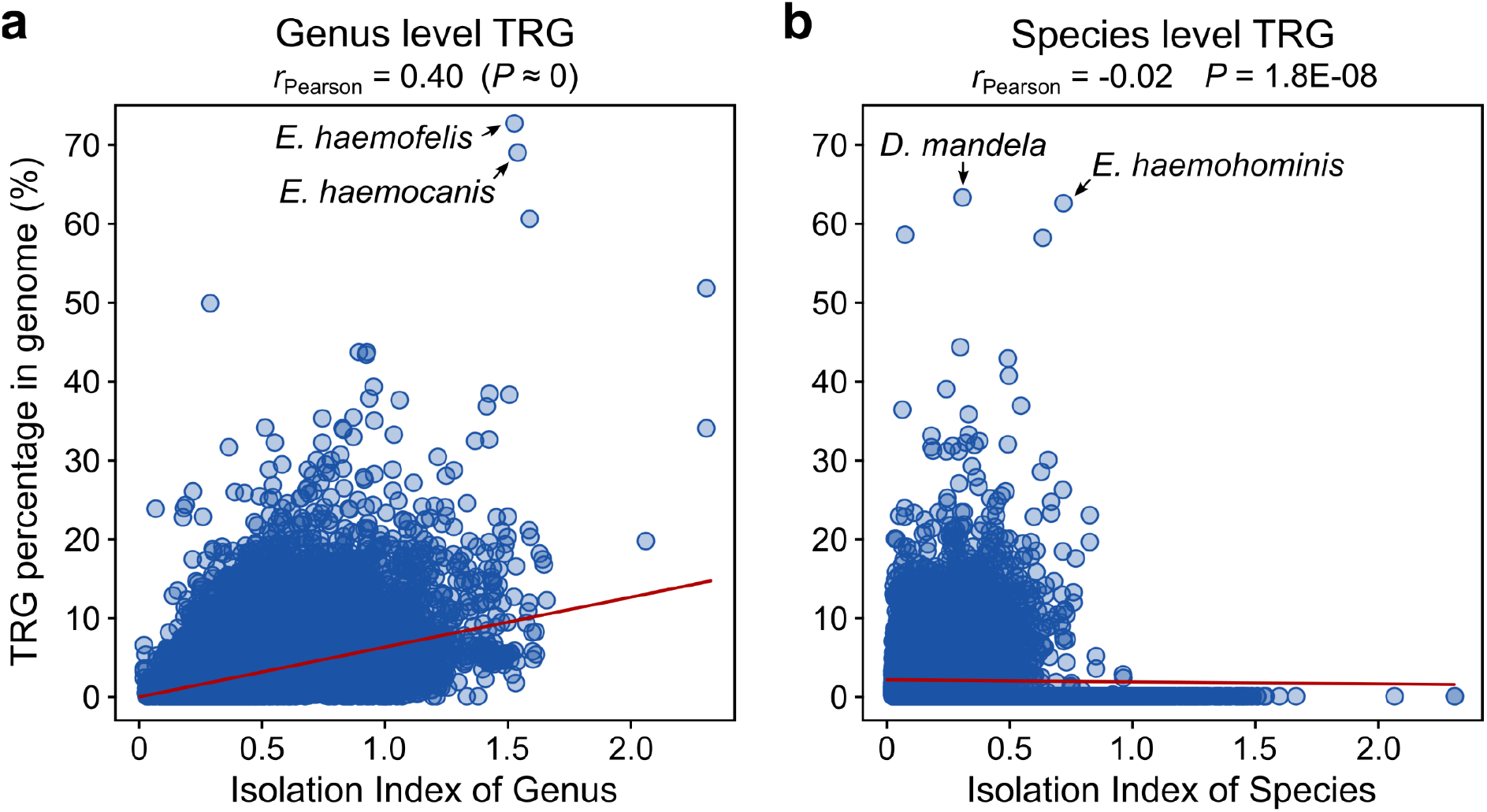
Correlation between the TRG number and isolation index of organisms (IIO) for **a**. genus- and **b**. species-specific genes.

The highest number of species-level TRGs was found in the genome of *Didemnitutus mandela* (NCBI assembly accession: GCA_002591725), a bacterial symbiont of marine tunicates. Despite the relatively low taxonomic isolation of *D. mandela* (IIO = 0.29), 63.3% of its genes (1,936 out of 3,060) were classified as species-specific (**Fig. 1b**), and an additional 13 (0.4%) were shared only within the *Didemnitutus* genus. The exceptionally high number of species-level TRGs is supported by a recent study of the *D. mandela* genome [30], which highlighted extremely high number of genes without detectable homologs, suggesting that the specific environment and distinct lifestyle of this bacterium resulted in this unusual accumulation of the species-specific genes.

Conversely, the highest number of genus-specific TRGs was identified in bacteria belonging to the *Eperythrozoon_B* genus. These bacteria are assigned to the *Mycoplasmoidaceae* family and are responsible for a rare, sporadic, non-contagious, blood-borne disease in ruminants [31]. Two species, namely *E. haemofelis* and *E. haemocanis*, contain 72.7% and 69.0% of genus-level TRGs, respectively (**Fig. 1a**). The number of species-specific TRGs is relatively low for these species, with a maximum of 5.2% in *E. haemofelis*. Interestingly, among the four species classified in this genus, one (*E. haemohominis*) contains a large proportion of species-specific genes (62.6%, **Fig. 1b**) and 4.5% of genus-specific genes.

We did not find any TR genes at the species or genus levels in the genomes of 12 species. Six of these species belong to the *Enterobacteriaceae_A* family and represent endosymbionts of aquatic leaf beetles (from *Donacia, Plateumaris*, and *Neohaemonia* genera). Four other bacterial species are classified as uncultured *Pelagibacteraceae* bacteria, and belong to the *Pelagibacter_A, MED727*, and *CACNRH01* genera. The remaining two species are symbionts of aphids (*Serratia symbiotica*) and blood sucking fly *Lipoptena cervi* (Candidatus *Arsenophonus lipoptenae*). Since the species isolation index IIO for these genomes varies greatly (between 0.33 and 1.00 for species and from 0.05 to 0.50 for genus) it is very difficult to assign the effect of these predictions to adequate taxonomic saturation.

### Properties of the TRG protein sequences

Since the protein sequences in the GTDB database lack annotation descriptions, we mapped the sequences of the predicted TRGs to the corresponding NCBI protein records. Due to missing protein sequence data for some NCBI genome assemblies, only 35% of the predicted TRGs could be mapped to the identical sequences deposited at NCBI. To gain insight into the functional annotations of these proteins, we used a word frequency strategy because the information is not available in any formal annotation system like Gene Ontology or KEGG. As expected, the most repeated descriptors for TRGs present in 49% TRG proteins include “hypothetical”, “uncharacterized”, “partial”, “unknown”, and “putative” (**Fig. 2**). The second layer of functional characterization includes general, non-specific descriptors (1%) such as “plasmid”, “domain-containing”, “conserved”, or “membrane”. Finally, the less frequent descriptors suggest TRGs may be involved in processes such as “regulation”, “transcription”, “export”, and “transport”. Notably, one of the middle scoring description words of the TRG records includes the keyword “orphan”, highlighting the taxonomically restricted nature of the proteins. Together, these annotations align well with the postulated role of TRGs in adaptation and specific organismal functions.

**Figure 2.**
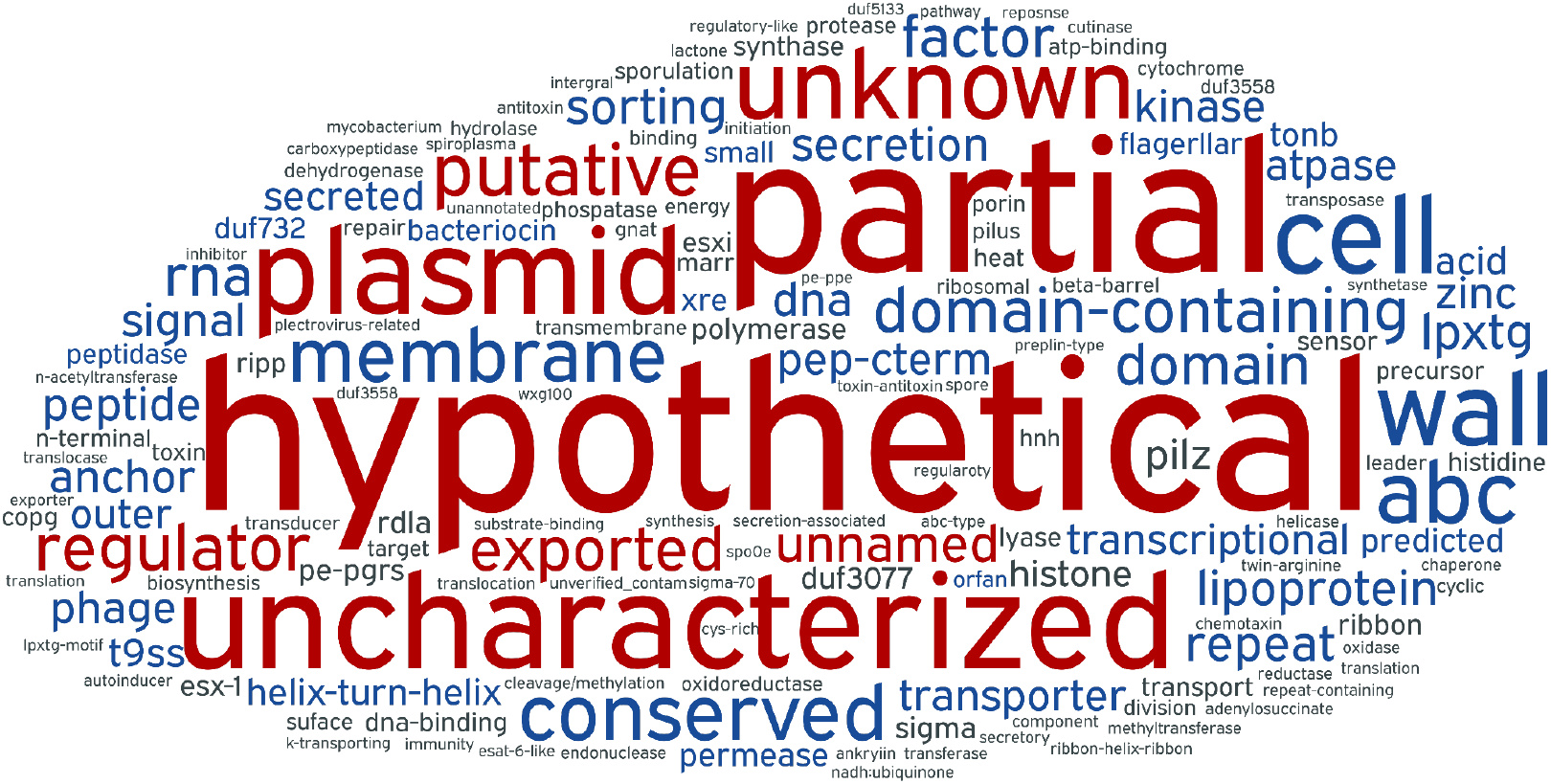
Word cloud extracted from NCBI Protein records of the TRG proteins. Common and meaningless words and characters were removed from the plot. Only words that appeared in at least 200 proteins are shown. The full list of words and their frequencies is available in the Supplementary File.

To further characterize the TRG proteins, we analyzed four common properties of these sequences: length, disorder and complexity levels, and aggregation potential. When compared to the properties of a randomly selected group of similar size from all non-TRG bacterial proteins, we observed that the TRG sequences are on average of smaller size (with a median of 67 and 87 for species- and genus-specific genes, respectively, compared to 281 for all other proteins; **Fig. 3a**) and are less complex (median ∼3.8 for TRGs versus ∼4.0 for background proteins; **Fig. 3b**), show higher disorder levels (median 0.16 and 0.18 for genus- and species-specific TRGs, respectively, and 0.06 for randomly selected proteins; **Fig. 3c**), and a slightly higher aggregation potential for genus-specific TRGs (median 4.9 versus 3.3 measured as background; **Fig. 3d**). The high aggregation potential of the genus-specific TRG sequences is a phenomenon noticed previously in *Bacillus* [21] and it may indicate that this group of genes has a unique evolutionary history.

**Figure 3.**
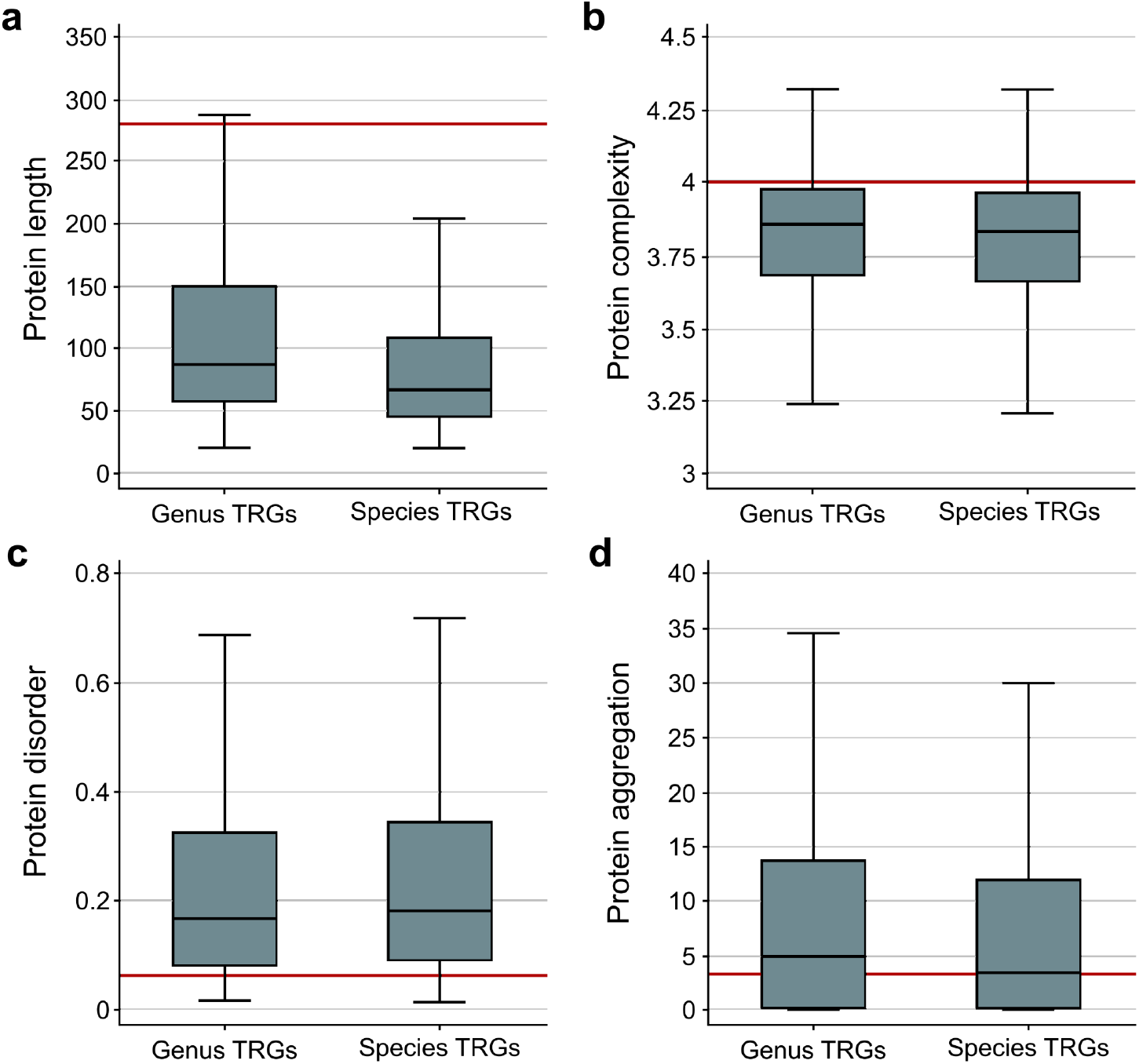
Comparison of protein sequence properties between the TR genes at the genus and species levels. Horizontal red line indicates the median value calculated from a random sample of the non-TRG bacterial proteins.

Finally, we provide insights into the evolution of the TR genes in bacteria by clustering their sequences based on similarity (see Methods). Although the majority of the TRG proteins (70%) do not form groups of evolutionary-related sequences (singleton clusters), we observed orthologs and paralogs on the genus level and paralogs on the species (single genome) level. The largest cluster was found in genomes belonging to the *Pseudomonas_E* genus and it covers 1,774 genus-level proteins. This cluster is composed of sequences coming from different species (orthologs) as well as encoded in the same genome (paralogs). Surprisingly, at the species level, we identified 283 sequences forming a group of homologous sequences in *Arsenophonus sp000757905*. It has to be noted that in this case, the sequences forming the cluster are very short, mainly 35 amino acids long.

### Database interface and functionality

The landing page of the TRGdb database provides the summary statistics concerning the number of predicted TRGs and their general characteristics (**Table 1**). The database interface offers simple and fast exploration capabilities, including browsing by taxonomy and searching by keyword and/or the TRG sequence.

The keyword search interface offers initial access to a table with all the bacterial species, allowing users to look for bacteria taxa of interest. For convenience, the search box features an autocomplete functionality that suggests terms matching the user query. Selecting a taxon name results in filtering the table rows. This procedure can be progressively applied to the previous search results.

The browse view provides a hierarchical exploration of the TR genes through bacterial taxonomy based on the GTDB classification. The interactive interface allows users to expand branches of the bacteria tree and view the number of genus and species level TRGs associated with each node. It begins with the superkingdom bacteria and selecting taxon names limits the range of selected records. Direct access to the list of genomes (species) is provided by choosing the “details” button (three dots icon). From that view, a user can access the genome-related statistics, including the list of the TR genes.

The TRG record is a collection of information about a selected sequence including external links to the information deposited at NCBI. However, since the TRG sequences are often poorly annotated and accompanied by limited information, the TRG records are constantly updated with new calculations using available tools and datasets. There are two ongoing projects that provide extra information about the quality of predictions and biological properties of the TRG sequences deposited in TRGdb. One project focuses on the high-quality annotation of the TRG sequences in the *Bacillus* genus [21], and these superior records are marked with a star in the database. The other project is a high-throughput effort to annotate the expression status of the TR genes, only considering high-quality predictions. This project uses publicly available RNA-Seq data sets that are specific to species with the TR genes and labels the TRG records with a positive expression status.

To enhance the search experience of TRGdb we also provide the BLAST feature to search for the TR genes based on sequence similarity. This option is useful for users who want to quickly check if their proteins of interest are classified as TRGs.

Finally, for customizable access to TRGdb, we provide an API interface allowing users to programmatically search and browse the database content and retrieve statistics on genome and the TRG records. The API interface of TRGdb has a dedicated help page listing all the functionalities. In addition, all the data that were used to construct TRGdb are also available for download including CSV files containing information on each bacterial genome and a TRG protein, and sequences in FASTA format of all TRG proteins at genus and species levels.

## Discussion

TRGdb represents the most comprehensive and up-to-date database of taxonomically restricted genes in bacteria. We aim to further identify and characterize these genes, and TRGdb serves as a valuable starting point for this journey. The database provides updated information on marker sequences in the context of the latest bacterial taxonomy, including analysis of their sequence properties and access to available annotations. Additionally, we continuously update TRGdb with data from RNA Seq analysis to improve the prediction quality and functional characterization of these genes.

The identification of TRGs in bacteria is a complex task that presents several challenges. One of the main challenges is the quality of the taxonomic classification. The classification of bacteria is a very dynamic field with a variety of different phylogenetic approaches (e.g., [32– 35]. Surprisingly, the advent of molecular and genomic data seems to have increased the variety of classifications rather than reducing the problem [36]. Since the TRG identification solely depends on the taxonomic classification of analyzed organisms, we decided to adopt the well-accepted and standardized solution provided in the GTDB database. Another challenge of the TRG identification is the computational cost of searching for TRGs. Even when limiting the available data to one representative genome for one species, the current data volume of available proteins is counted in hundreds of millions. The well-accepted solution to search for sequence similarity during the TRG annotation is BLAST. However, even when executed on high computational power clusters, searches with BLAST would take months of computational time. To ensure the up-to-date status of our database, we have designed a simplified mode of the TRG annotation pipeline. The predictions are processed in three steps, where the first is designed to limit the search space for the application of BLAST search. In this way, at the current data volume, we can recalculate the whole database within ten days and thus keep up with the annual GTDB updates.

The increasing number of TRGs is one of the greatest surprises in the field of bacterial genome sequencing [5]. We also observe this trend in our database – the comparison of the first publicly available release of the TRGdb database with the previous version (based on GTDB release 202 from 27 April 2021) shows that the number of predicted TRG sequences has increased by 25% (the current 8.5 versus previous 6.8 million sequences). It has to be noted, however, that the previous release of the GTDB database covered only 45,555 species, while the current version contains 62,291 species, and the mean number of TRGs per species between TRGdb releases seems constant (approximately 4%). This comparison demonstrates the dynamic nature of TRGs and the importance of keeping a regularly updated database with the most current information on bacterial taxonomically restricted genes.

## Acknowledgments

The computations were performed at the Poznan Supercomputing and Networking Center (PSNC) within grant numbers 312 and 528.

## Data availability

The TRGdb database along with documentation is available at http://combio.pl/trgdb

## Author contribution

AZ performed the calculations, analyzed the data, and implemented the database and web interface; WD implemented the API interface and provided data from previous database releases; WMK designed the study, analyzed data, and wrote the manuscript.

